# Enhancing Germination and Assessing the Longevity of Native Plants’ Seeds with Dormancy through Different Priming Techniques

**DOI:** 10.64898/2026.10.30.763245

**Authors:** Majda Khalil Suleiman, Shabbir Ahmad Shahid, Narayana Ramachandra Bhat, Sheena Jacob, Mini Thiruthath Sivadasan, Rini Rachel Thomas, M. Anisul Islam

## Abstract

*Vachellia pachyceras* and *Convolvulus oxyphyllus* are native species of arid and semi-arid desert regions, including Kuwait. They are inherently adapted to extreme conditions characterized by high temperature, prolonged drought and limited rainfall. They contribute significantly, to restore the ecosystem by providing food and shelter for wildlife. Despite their ecological importance, seed germination in both species is constrained by dormancy, and natural recruitment of seedlings is limited by narrow and unpredictable rainfall windows, severely restricting opportunities for establishment. In this context, seed priming is an effective pre-sowing technique that enables uniform and rapid germination within the short window of favorable conditions. This study aims to identify the optimal seed-priming techniques for these two keystone native species and to assess the effect of priming on seed longevity. The seeds were scarified to remove the physical dormancy prior to priming treatments. Relative seed longevity was measured for primed seeds through artificial ageing and analyzed using probit analysis and analysis of variance. Results showed that hormonal priming [(mechanical scarification + 100 ppm gibberellic acid (GA_3_)] for six hours and hydropriming (mechanical scarification + soaking in water) for 8 hours (h) were effective treatments among the tested priming treatments for *V. pachyceras.* The estimated p50 value (i.e., number of days to lose seed viability to 50%) was 22 days for hormonal priming and 21 days for hydropriming. Mechanical scarification followed by one and half hour of hydropriming was the most effective in *C. oxyphyllus* (p50 = 74 days). Based on these results, the longevity of *V. pachyceras* and *C. oxyphyllus* could be considered ‘medium’. Further studies are needed to determine the effectiveness of seed priming on additional ecologically and economically important native species to achieve success in revegetation programs.

**Graphic Abstract:** 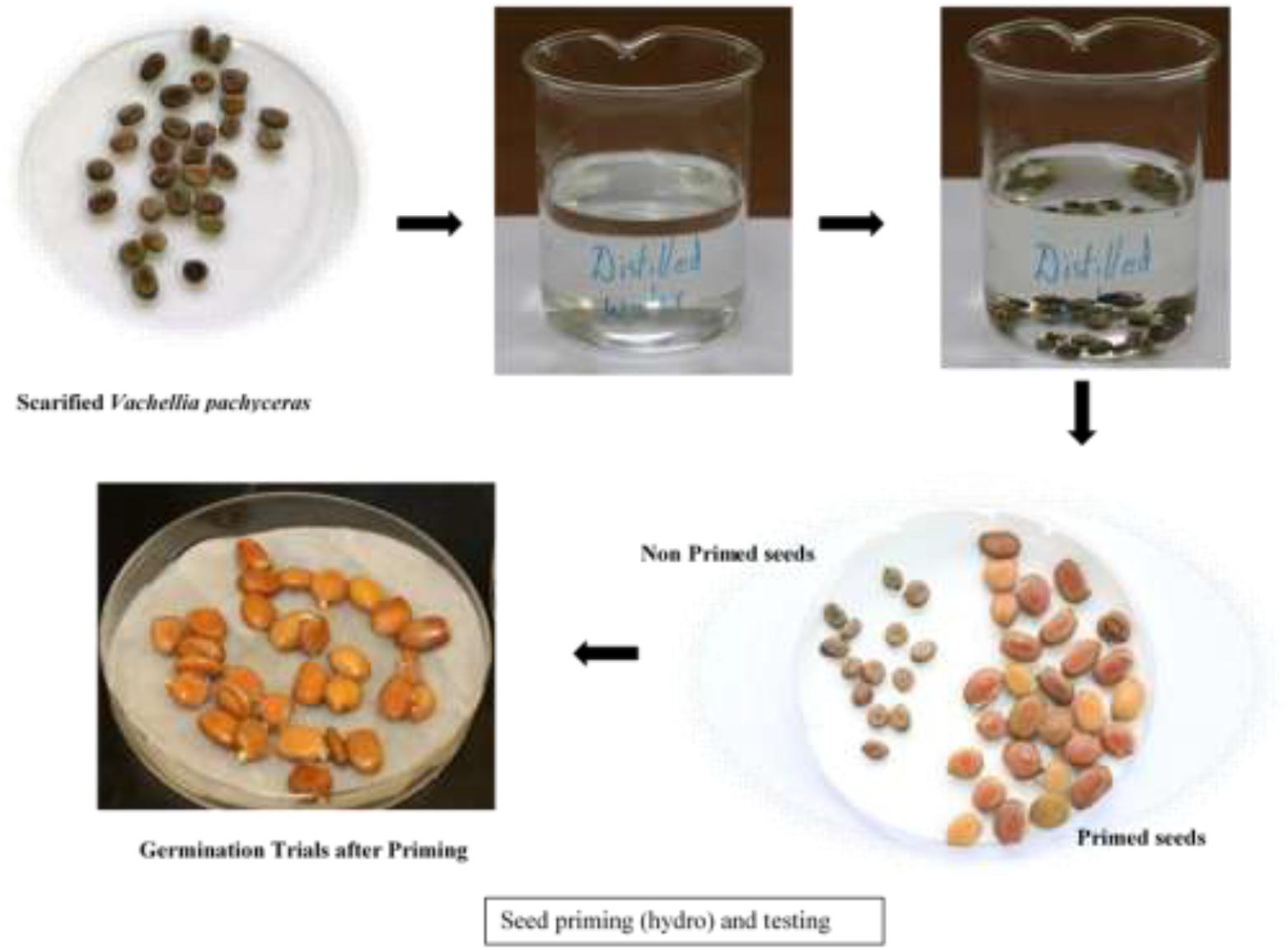

**Highlights:**

- Seed priming with species-specific priming treatments can help in securing uniform, faster, and higher germination in selected native desert plants.
- Seed enhancement treatments like seed priming ensure better survival and seedling establishment, which is critical for revegetation programs, as natural seed production of native seeds in arid conditions is minimal.
- The study ranks the priming treatments for *Vachellia pachyceras* and *Convolvulus oxyphyllus* and recommends hydropriming (Mechanical Scarification + soaking in water for 8 h) for *Vachellia pachyceras*, given the cost and ease of use.
- Mechanical scarification followed by 1.5 h of hydropriming was the most effective priming treatment for *Convolvulus oxyphyllus* seeds.
- These results suggest that primed seeds of these two species have medium longevity and can be stored in seed banks and used for seed-based restoration programs.

## 1. Introduction

In the hyper-arid regions of the world where aridity index (rainfall/evaporation) is extremely low, ecological restoration of damaged ecosystems is critically important. Kuwait is one such arid region characterized by extreme temperatures during summer and, winter, poor sandy soil, erratic and low rainfall [1]. Both anthropogenic disturbances and natural effects have contributed to a rapid decline in native vegetation, reducing plant cover to approximately 10% of total area over the past decades [2–5].

Among the 374 native species recorded in Kuwait, *Vachellia pachyceras O. Schwartz (Family: Fabaceae Sub family: Mimosoidae)* is the only native tree species and currently represented by a single naturally occurring mature individual in Sabah Al-Ahmad Natural Reserve. *Convolvulus oxyphyllus Boiss; (Family Convolvulaceae)* is a perennial native woody shrub, and one of the 15 key desert plants of Kuwait. It is associated with Rhanteriatum vegetation community and the total area covered by this community is 2%. However, specific population data for C. oxyphyllus in Kuwait remain limited and it is considered part of the vulnerable ecosystem. Seeds of both these species exhibit physical dormancy due to hard impermeable seed coat

As an adaptation to arid conditions, many of the desert species possess inherent dormancy mechanisms. Due to unfavorable climatic conditions, the natural seedling recruitment is extremely limited with the probability of a seed developing into a mature plant estimated to be 10% [6–7]. The rare event of germination and establishment leading to significant natural recruitment might take decades, and therefore, deliberate attempts of timely revegetation are essential [8]. In Kuwait, native plant species have recently been incorporated into revegetation programs [9–10]. However, multiple biotic and abiotic ecological barriers like seed dormancy, extreme temperatures, erratic rainfall, poor soil quality, grazing continue to hinder restoration success, especially during the early seedling development stages [11]. Seedling emergence and survival during the narrow and unpredictable rainfall windows remain a key limiting factor for plant establishment.

In this context, seed enhancement technologies offer promising solutions to overcome major barriers of restoration. These technologies involve physical and biochemical manipulation to seeds, to enhance germination percentage and rate, seedling emergence, and early seedling establishment and growth [12–13]. Pretreated seeds have been shown to germinate quickly under very low soil moisture conditions and make the most efficient use of limited and highly variable soil moisture levels available under arid conditions [14–15].

Seed priming is a widely used seed enhancement technique which involves controlled hydration of seeds to initiate some of the physiological processes of germination without allowing radicle emergence followed by re-drying prior to sowing [16–18]. This pre-treatment provides “head start” enhancing germination, seedling vigor, and tolerance to harsh conditions like drought and salinity. Priming methods range from simple hydropriming (using water) to osmopriming (using osmotic solutions), hormonal priming [19], biopriming (using beneficial microbes), and chemical priming [20].

In the hydropriming technique, seeds are soaked in water before sowing, which may be followed by drying [21]. In contrast, osmopriming involves soaking seeds for a specific period in osmoticum solutions that allow controlled hydration such as Polyethylene Glycol (PEG) at water potentials below 0 MPa, followed by drying [21] PEG, typically with a molecular weight of 6000 to 8000, is a most commonly used osmotic agent for osmopriming. By increasing the osmotic pressure of the solution, PEG regulates water availability and slows the rate of seed hydration, thereby reducing the cellular damage associated with rapid water uptake by dry seeds [22]. Furthermore, due to its high molecular weight, PEG is readily absorbed by seeds, minimising the risk of toxicity during the priming process. In hormonal priming, seeds are treated with optimal concentration of phytohormones like Indole-3-acetic acid (IAA), Abscisic Acid (ABA) and Gibberellins (GA) for an appropriate duration, and are subsequently air-dried [19]. Priming the seeds with GA_3_ promotes germination by mobilizing the stored reserves and stimulating embryo growth [23]. Combination priming involves combining two or more priming methods as a seed enhancement strategy. However, the effectiveness of these treatments depends on species-specific responses, and therefore the concentration of priming agents and duration of the treatment must be standardized for each species.

Despite growing recognition of seed enhancement technologies in restoration ecology, their application remains limited, particularly for native plant species. Pedrini et al. [21] reported only 14 peer reviewed studies on seed enhancement in native species published between 2006 to 2020. Notably, no studies are available on seed priming in *Vachellia pachyceras* and *Convolvulus oxyphyllus*, both of which are critical for restoring degraded arid landscapes in Kuwait. Most existing research is focused on agricultural crops such as Wheat [24] cereals and horticultural crops [25] highlighting a significant research gap.

Generally, seeds of some species can be stored under certain conditions, which allow them to maintain their viability for a long time, but their longevity can vary greatly between species [26–27]. The term seed longevity refers to the time period during which the seed remains viable and capable of germination under favourable conditions [28]. Hydration and subsequent drying during seed priming may affect seed longevity, as moisture and temperature are key determinants of seed lifespan [29–30]. Consequently, priming induced changes in seed can influence their viability and storage behavior.

Knowledge of longevity of primed seeds is essential if it is intended to be carried out commercially to have an idea of period of storage of the primed seeds. Direct measurement of seed longevity under storage (ideally -20 °C) conditions and in real time can take years as it may take decades for the decline of seed viability [26]. Therefore, as an alternative, seed ageing is accelerated (artificial ageing) by storing the seeds at relatively high temperature (45 °C) and moist conditions [31–34]. Originally developed by Delouche [35], these tests are widely used for quality control by seed companies, and provides reliable information to estimate storage potential of a particular batch of seeds. Artificial ageing has also been used to evaluate performance of seedlings under field conditions and to differentiate the physiological attributes of seed lots with similar emergence behavior [36]. It has been reported that if the quality of seeds deteriorates during the accelerated ageing process and conditions, their ability to germinate will also be lost eventually [37]. Furthermore, studies [38–39] have found a correlation between natural and artificial ageing of seeds in three woody perennials and the mechanisms that accelerate the deterioration to be same in both natural and artificial ageing – differing only at speed of the process. Therefore, by conducting controlled ageing experiments, one can rank seed longevity of a species relative to other species by estimating the time taken by 50% of the seeds to lose viability [31].

Although the mechanism underlying the effects of seed priming on germination and seed longevity in different species are yet to be understood; however, it is known to enhance germination percentage, and uniformity by mechanical weakening of endosperm and development of endo-β mannanase activity [40]. However, the efficacy of different seed priming on native desert plant species remains largely unexplored.

This study aims to standardize different seed priming techniques including hydropriming, hormonal priming, osmopriming, and combinational treatments to enhance germination in two important native species (*V. pachyceras* and *C. oxyphyllus*). Additionally, it evaluates the effects of priming on seed longevity using artificial ageing tests (*V. pachyceras* and *C. oxyphyllus*). It is hypothesized that seed priming will have a positive effect on germination rate and germination percentage, thereby contributing to more effective restoration of degraded desert ecosystems. The ultimate objective is to identify optimal priming treatments suitable for large-scale revegetation efforts in Kuwait.

## 2. Materials and Methods

### 2.1. Seed Sources

Seeds of *V.pachyceras* were procured from Wadi al-Faisah, Saudi arabia in in 2017, and the seeds of *C. oxyphyllus* were collected from Sabhan area in Kuwait in 2017. Seeds were stored in refrigerator at 4 °C prior to the initiation of the priming studies. The priming and longevity studies were conducted during 2018-2019.

### 2.2. Determining Optimal Priming Treatments

Imbibition and germination process [41] of seeds includes three phases: an initial rapid water uptake (Phase 1), a slower phase of water uptake, during which metabolic reactivation occurs (Phase 2) and radicle protrusion (Phase 3). During the priming treatment, the seeds are exposed to specific priming agents and maintained in phase 2, preventing progression to radicle protrusion. Prior to conducting the priming treatments, it is essential to standardize both the concentration of the priming agent and the duration for each species, as responses vary significantly among species. *V. pachyceras* and *C. oxyphyllus* seeds were either scarified using sand paper (mechanical scarification) or with acid (Concentrated Sulphuric acid for 30 min), prior to standardization procedures to facilitate water imbibition which is necessary to initiate the germination process and to overcome physical dormancy [42–44]. For acid scarification, the seeds are soaked in concentrated Sulphuric acid for the pre-determined time (30 min), followed by thorough rinsing under cool running water for 10-15 minutes till it is free of acid.

#### 2.2.1. Hydropriming

##### Standardization of Hydropriming

In order to determine the priming duration, scarified (acid and mechanical) and control seeds were soaked in water, and the weight gain was recorded as mentioned below to prepare the imbibition curve. Five replications of 10 seeds for each species were placed on moistened filter paper in Petri dishes at 25 °C and every hour they were patted dry with tissue paper, and the total seed weight of all the seeds in each replicate was recorded [45]. The experiments were terminated at radicle protrusion (1 mm length). Percentage weight gain was calculated as described by Jeyasuriya et al. [46], and the data were used for the preparation of the imbibition curve.

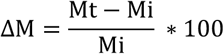

where, ΔM = increase in mass; Mt = seed weight after water uptake for time t; and Mi = initial seed weight.

Optimal hydropriming treatment durations were determined from the imbibition curve when water uptake in the first linear phase declined or remained the same. The selected treatment durations are detailed in Table. 1

Hydropriming Treatments: Seeds were hydro-primed as following: three replications of 30 seeds per treatment were primed (Table 1) and kept for germination. Control seeds were not subjected to priming. Seeds of *V. pachyceras* and *C. oxyphyllus* were placed in Jiffy pots and Petri dishes respectively in desert soil medium (non-saline ECe 0.3 mS/cm; sandy texture and slightly alkaline reaction (pHs 7.5). Jiffy pots were filled with desert soil and the *V. pachyceras* seeds were sown at 1 cm depth. For *C. oxyphyllus*, the petri dishes were filled with desert soil at 1 cm depth and the seeds were placed on the surface, as the seeds of *C. oxyphyllus* are small in size. The primed seeds were maintained under laboratory conditions at 25 °C and the experiment was continued for 30 days or more depending on the species. Seeds were closely monitored for germination each day, and germination data was recorded. Radicle protrusion of 1 mm was considered as germination. Based on the germination percentage, rate, and vigor, the best priming treatment was selected for longevity experiments and the results are presented in section 2.2.4.

**Table 1.** Hydropriming Hours for Each Species Based on the Hydration/Imbibition Curve.

| *Species* | *Scarification Treatment* | *Hydropriming Hours* |
|---|---|---|
| *Vachellia pachyceras* | Non-scarified seeds | 0 (Control), 8, 12, and 16 h |
|  | Acid scarified | 0, 4, 8, and 12 h |
|  | Mechanically scarified | 0, 4, 8, and 12 h |
| *Convolvulus oxyphyllus* | Non-scarified seeds | 0 (Control) , 2, 4, and 6 h |
|  | Acid scarified | 0, 45 min, 1h 30 min, 2 h |
|  | Mechanically scarified | 0, 45 min, 1h 30 min, 2 h |

#### 2.2.2. Hormonal-Priming

##### Standardization of Hormonal Priming

In order to determine the hormonal priming duration, scarified and control seeds were soaked in GA_3_ (prepared from GA_3_ Sigma CAS No. 77-06-5), and the weight gain was recorded as mentioned below to prepare the imbibition curve. A set of ten seeds with three replications were treated with GA_3_ (100 ppm GA_3_ for *V. pachyceras* and 750 ppm GA_3_ for *C. oxyphyllus*) and were placed in Petri dishes lined with moistened filter paper at 25 °C. Concentration of GA_3_ for *V. pachyceras* was finalized based on the unpublished experimental observations at KISR and for *C. oxyphyllus*, the concentration was based on the results by [42]. After 1 h, the seeds were patted dry with tissue paper and the total weight of all the seeds in each replicate was recorded using a digital weighing balance. The same procedure was repeated after regular intervals. The experiments were terminated at radicle protrusion (1 mm length). Water uptake was determined gravimetrically, and the percentage weight gain was calculated as mentioned in hydropriming experiments. From the imbibition curve for seeds treated with GA_3_, hormonal priming hours were determined (Table 2).

**Table 2.** Hormonal Priming Hours for Each Species.

| *Species* | *Treatment* | *Duration of Hormonal Priming* |
|---|---|---|
| *Vachellia pachyceras* | Non-scarified seeds treated with 100 ppm GA<sub>3</sub> | 0, 6, 12, and 24 h |
|  | Mechanically scarified seeds treated with 100 ppm GA<sub>3</sub> | 0, 6, 12, and 24 h |
|  | Acid scarified seeds treated with 100 ppm GA<sub>3</sub> | 0, 6, 12, and 24 h |
| *Convolvulus oxyphyllus* | Non-scarified seeds treated with 750 ppm GA<sub>3</sub> | 0, 30, 60, and 90 min |
|  | Mechanically scarified seeds treated with 750 ppm GA<sub>3</sub> | 0, 30, 60, and 90 min |
|  | Acid scarified seeds treated with 750 ppm GA<sub>3</sub> | 0, 30, 60, and 90 min |

Hormonal Priming Treatments: After determining the optimum duration of hormonal priming, three replications of 30 seeds per treatment were primed (Table 2) and kept for germination. Control seeds were not primed. Seeds of *V. pachyceras* and *C. oxyphyllus* were placed in Jiffy pots and Petri dishes respectively in desert soil medium. The primed seeds were maintained under laboratory conditions at 25 °C and the experiment was continued for 30 days or more depending on the species. Seeds were closely monitored for germination each day, and observations were recorded. Based on the germination percentage, rate, and vigor, the best hormonal priming treatment was selected for longevity treatments and presented in section 2.2.4.

#### 2.2.3. Osmopriming

##### Standardization of osmopriming

To determine the optimal osmopriming treatment for each of the two species, mechanically scarified seeds were subjected to different osmotic concentration gradients of polyethylene glycol (PEG 8000) [0, -0.1, -0.4, -0.8, -1, - 1.5 and -2.0 MPa] at room temperature (25 °C). Each experiment was replicated three times with 20 seeds in each replication. PEG 8000 was mixed with deionized water to obtain different concentrations of osmotic potentials as per Hardegree and Emmerich [47]. Deionized water, which had an osmotic potential of 0.0 MPa, was used as control. Seeds were placed in Petri dishes with filter paper and soaked with 15 ml of PEG solutions. Germination of seeds was monitored for 14 days and the optimal priming concentration (least negative water potential for that species) was considered to be the one in which seeds did not germinate [48–50]. The least negative water potential marks the boundary where external water availability is insufficient to complete germination. A full two-week treatment window of no germination ensures all seeds attain a uniform pre-germinative readiness state. The optimal priming potential for *V. pachyceras and C. oxyphyllus* was -1.5 MPa.

Osmopriming: After identifying the optimal osmopriming treatment, seeds of each species were treated with the aforementioned osmotic potential (-1.5MPa) for 2, 4, 6 and 8 days at 25 °C. Then the seeds were washed thoroughly, spread on a tissue paper and dried at 25 °C under lab conditions prior to germination tests, while control seeds were not osmoprimed. The experiments were conducted with three replications with 30 seeds each. The primed seeds were maintained under laboratory conditions at 25 °C and the experiment was continued for 30 days or more depending on the species. Seeds were closely monitored each day, and the observations were recorded. After the germination and vigor tests were conducted, best osmopriming treatments (osmotic concentration and duration of treatment) for each species were optimized and the results are presented in section 2.2.4.

#### 2.2.4. Combination priming (Combination of hydro and osmo)

*Standardization of treatments:* Seeds of each species were subjected to the best hydro-priming treatments identified from the previous experiments (Table 3), dried, and then again osmoprimed (as per the osmopriming protocol described above) in 0, -0.1, -0.4, -0.8, -1.0, -1.5, and -2.0 MPa PEG 8000 solutions at 25 °C to evaluate their germination for 14 days. The combination of hydropriming and osmopriming treatments, which did not result in germination for 14 days, was selected for the combination priming. Each experiment was replicated three times with 20 seeds in each replication.

**Table 3.**
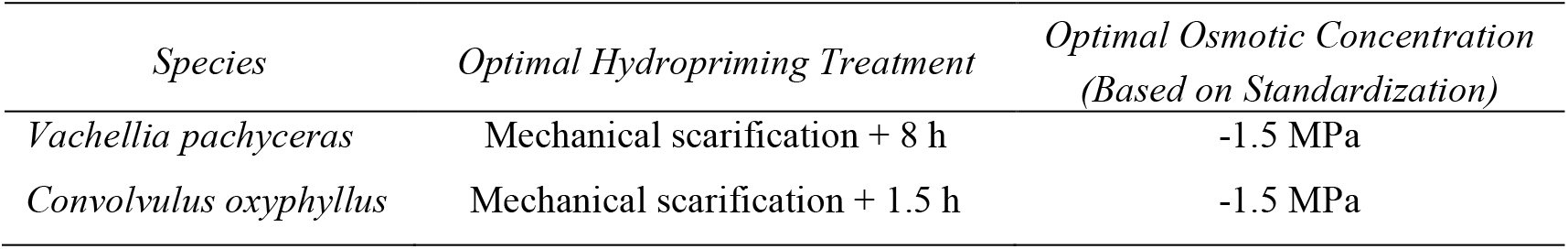
Optimal Hydropriming and Osmopriming Treatments Used as Baseline for Combination Priming.

##### Combination priming

The hydroprimed seeds were dried and subjected to the selected osmopriming solutions, which were derived from the results of the aforementioned standardization procedure for each species for 2, 4, 6, and 8 days at 25 °C, to determine their responses. Then the seeds were washed and dried at room temperature (25 °C) and kept for germination under laboratory conditions at 25 °C and the experiment was continued for 30 days or more depending on the species. Control seeds were not subjected to priming treatments. There were three replications with 30 seeds in each replication. Seeds were closely monitored each day, and the observations were recorded.

After conducting the combined priming treatments, observations on germination percentage, time to 50% germination, and vigor test were recorded for the assessment of optimal priming treatment for each species.

After selecting the best priming treatment from hydropriming, osmopriming, hormonal priming, and combination priming experiments (Table 4), the seeds were subjected to optimal priming treatments followed by artificial ageing to evaluate the effect of seed priming on seed longevity. The results of the experiments for selection of best priming treatment is provided in the supplementary material.

**Table 4.** Optimal Seed Enhancement Priming Treatments Selected for Longevity Experiments.

| **Species** | **Optimal Priming Treatments** |  |  |  |
|---|---|---|---|---|
|  | **Hydropriming** | **Hormonal Priming** | **Osmopriming** | **Combination Priming** |
| *Vachellia pachyceras* | MS + 8 h | MS + 100 ppm GA<sub>3</sub> for 6 h | MS + 2 days in -1.5 MPa PEG | MS hydro 8 h + 4 days in -1.5 MPa PEG |
| *Convolvulus oxyphyllus* | MS +1.5 h | MS + 750 ppm GA<sub>3</sub> for 30 min | MS + 2 days in -1.5 MPa PEG | MS hydro 1.5 h + 2 days in -1.5MPa PEG |

MS: Mechanical scarification; GA<sub>3</sub>: Gibberellic acid; PEG 8000: Polyethylene Glycol 8000.

##### Longevity experiments

Standard ageing procedures [27, 31, 34 and 51] were used for conducting the longevity trials of the primed seeds. In open glass Petri dishes (52 × 12 mm), 18 samples of 50 primed seeds each were placed in a single layer on the incubation stand (Fisher brand incubation tray in polypropylene blue, Fisher Scientific Ltd., UK). This was placed over a non-saturated solution of LiCl (anhydrous, laboratory reagent grade, Fisher Scientific UK Ltd, Leicester, UK) in airtight electrical enclosure boxes with clear lid (300 × 300 × 130 mm, Ensto UK Ltd, Southampton, UK) and was rehydrated at 47% relative humidity (RH) (385 g of LiCl in one liter of distilled water) and 20 °C temperature. At the end of the rehydration period of 14 days, the seed equilibrium relative humidity (eRH) of the equilibrated seeds was checked using a rotronic hygrometer (AW-DIO sensor with HygroPalm 3 display unit, Rotronic Instruments UK Ltd, Crawley, UK) to confirm that the seeds had attained equilibrium.

The rehydrated primed seeds were then transferred to the incubation trays in the second electrical enclosure box over a non-saturated solution of LiCl (ageing solution) at 60% RH (300 g of LiCl in one liter of distilled water) and placed in an incubator at 45 °C in dark. The RH of the LiCl solution was checked periodically using tiny tag View 2 -TV 4050 (Gemini Data Loggers, UK), and the necessary adjustments were made by adding the required amount of distilled water to the solution in the enclosure box.

One sample of 50 seeds of *V. pachyceras* was withdrawn after 0, 1, 2, 3, 4, 5, 6, 7, 8, 9, 10, 20, 30, 60, 120, 150 days for each ageing experiment while same number of *C. oxyphyllus* seeds were withdrawn after 0, 1, 2, 3, 4, 5, 6, 7, 8, 9, 10, 20, 30, 60, 120, 150, 210, and 240 days. From the project team’s previous observations, it was presumed that the longevity of *C. oxyphyllus* would be more, and hence, 210 and 240 days were included with the ageing experiments for this species. As per the planned retrieval days, the seeds of both *V. pachyceras,* and *C. oxyphyllus* were retrieved and incubated for germination in soil medium at 25 °C. At each retrieval date, a single bulk sample of 50 seeds was removed and subsequently divided into five sub samples for the germination assay. The germination data were recorded daily and seeds with a radicle growth of ≥2 mm were considered to be germinated.

##### Recording of observations

Germination was recorded daily and the biomass was recorded at one month after sowing. Germination percentage and time to 50% germination were recorded [52–53]. The seedling vigor was evaluated based on the speed with which germination was achieved. The number of abnormal seedlings was also recorded, which was considered to calculate the germination percentage using the following equation.

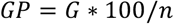

where *GP* = germination percentage; *G* = number of normal seedlings; and *n* = number of seeds sown [54].

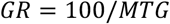

where *GR* = germination rate (in days) and *MTG* = mean time to germination [55].

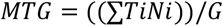

where *Ti* = the day of germination (from planting); *Ni* = number of seeds germinated at *Ti;* and *G* = total number of germinated seeds [56–57].

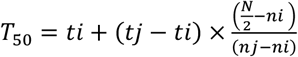

where *T*_50_ is the time (no. of days) to 50% germination; *N* is the final number of germination; and *ni* and *nj* are cumulative number of seeds germinated by adjacent counts at times *ti* and *tj* and *ni*<*N*/2<*nj* [52, 58].

Seedling vigor index (*SVI*) was calculated at the end of the germination test using the following equation [59]:

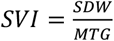

where *SDW* is seedling dry weight and *MTG* is mean time to germination. Seedlings were weighed and dried in an oven at 75 °C for 24 h to determine the dry weight of seedlings.

##### Data analysis

For longevity studies, a generalized linear model with binomial error and probit link function was fitted to the survival data (number of seeds germinating, number of seeds sown) in ®GenStat 21^st^ Edition (VSN International Ltd., Oxford, UK) with the storage period as an explanatory variable and both replicates and treatments as factors, thereby fitting the viability equation [59] as follows:

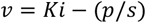

where v is the viability (in normal equivalent deviates, NED) of the seed lot after p days in storage, Ki is the initial viability (NED) of the seed lot, and s is the time (days) for viability to fall by 1 NED (i.e., the standard deviation of the normal distribution of seed deaths over time). From the probit analysis, p50 value for each replication in each treatment was derived and these values were analyzed using ANOVA and when treatment was significant, Duncan’s multiple range test was done to ascertain the statistical significance among them.

For the longevity experiments, germination data (germination rate, seedling vigor, germination percentage, and time to 50% germination) were analyzed by ANOVA using Genstat software (21st Edition; VSN International Ltd., Oxford, UK) to assess the effects of seed treatments on germination. When the main effect of treatment was significant, treatment means were compared using Duncan’s multiple range test method and significant differences among treatments were ascertained with alpha level of 0.05.

The germination percentage data were tested for the homogeneity of variance using Bartlett’s test. The square root of percentage of germination was arcsine transformed prior to the analysis if the data were non homogenous, but while presenting the data, non-transformed means are used as per the procedures of Mucunguzi and Oryem-Origa [55] and Bishop [60]. However, limitation of the analysis is that single bulk sample was retrieved at the selected time and the subsamples were treated as replicates. Hence it might result in overestimated degrees of freedom.

## 3. Results

### V. pachyceras

Data of probit analysis and subsequent ANOVA using the Genstat software revealed that p50 (i.e., number of days to lose seed viability to 50%) value of seeds that were hormone primed (22 days) and hydroprimed (21 days) were significantly different (*p* < 0.001) than the osmoprimed and combination primed seeds (Figure 1 and Table 5).

**Figure 1.**
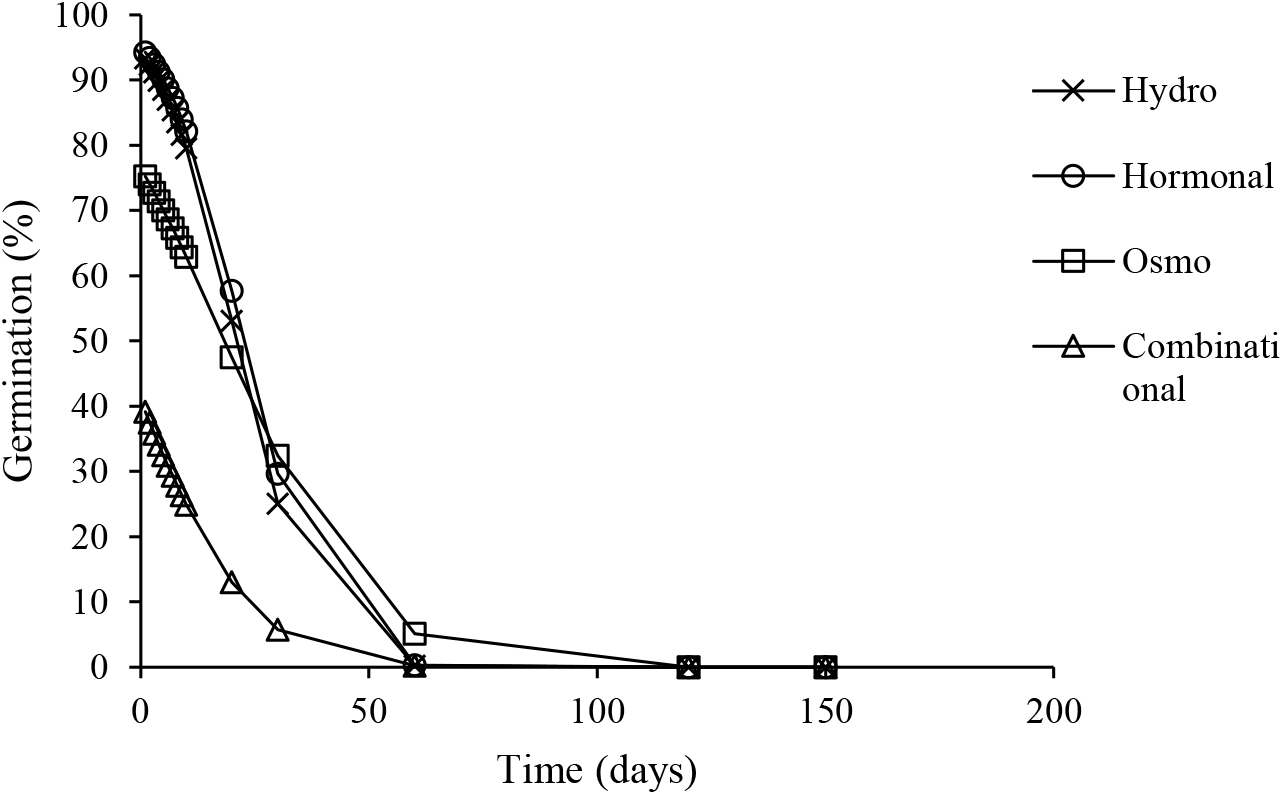
Survival curves fitted using a generalized linear model with binomial error and probit link function using ®GenStat 21^st^ Edition (VSN International Ltd., Oxford, UK) for ageing experiments in hydroprimed, hormone primed, osmoprimed, and combination (hydro + osmo) primed *Vachellia pachyceras* seeds.

**Table 5.** Survival Curve Parameters for Primed *Vachellia pachyceras* and *Convolvulus oxyphyllus* Seeds Determined Using a Generalized Linear Model with Binomial Error and Probit Link Function.

| **Treatment** | **Ki (NED)** | **SE** | **Sigma (1/days)** | **SE** | **Sigma (Days)** | **p50 (Days)** |
|---|---|---|---|---|---|---|
| ***Vachellia pachyceras*** |  |  |  |  |  |  |
| Hydropriming | 1.574 | 0.09 | -0.07487 | 0.00673 | 13.4 | 21 ab |
| Hormonal priming | 1.6494 | 0.09 | -0.0728 | 0.00651 | 13.7 | 22 a |
| Osmopriming | 0.7213 | 0.07 | -0.03923 | 0.00428 | 25.5 | 18 b |
| Combination priming | -0.2299* | 0.07 | -0.04483 | 0.00757 | 22.3 | * c |
| ***Convolvulus oxyphyllus*** |  |  |  |  |  |  |
| Hydropriming | 0.8515 | 0.0631 | -0.011428 | 0.00088 | 87.5 | 74 a |
| Hormonal priming | 0.1437 | 0.054 | -0.0102 | 0.00101 | 98.0 | 14 b |
| Osmopriming | 0.2205 | 0.0521 | -0.00999 | 0.00095 | 100.1 | 20 b |
| Combination priming | 0.2412 | 0.054 | -0.01233 | 0.0012 | 81.1 | 20 b |

Survival curves fitted using a generalized linear model with binomial error and probit link function using ®GenStat 21st Edition (VSN International Ltd., Oxford, UK) for ageing experiments in hydroprimed, hormone primed, osmoprimed, and combination (hydro + osmo) primed *Vachellia pachyceras* and *Convolvulus oxyphyllus* seeds. Values of p50 were compared using Duncan’s multiple range test method. There were five replications(subsamples) with 10 seeds in each replication. The mean values with the same letter are not significantly different at P <0.05
\*The viability of *V. pachyceras* was <50 at outset, by the time, combination priming was completed.

Ageing duration also affected the germination parameters of hydroprimed seeds. Increased ageing duration reduced germination percentage (Figure 2), germination rate, SVI, while increasing the time to 50% germination (Table 6). For instance, germination rate was the highest after two days of ageing treatment and gradually declined thereafter. No germination was observed when the seeds were aged more than 30 days (Figure 2A). The highest SVI was recorded in seeds with no ageing treatment (control) and in seeds aged for two days (Table 6). Consistent with the trend in germination rate, SVI decreased as ageing duration increased up to 30 days. For hydroprimed seeds, the highest germination percentage was observed at zero (0) days of ageing and declined steadily as ageing progressed to 30 days (Figure 2A).

**Figure 2.**
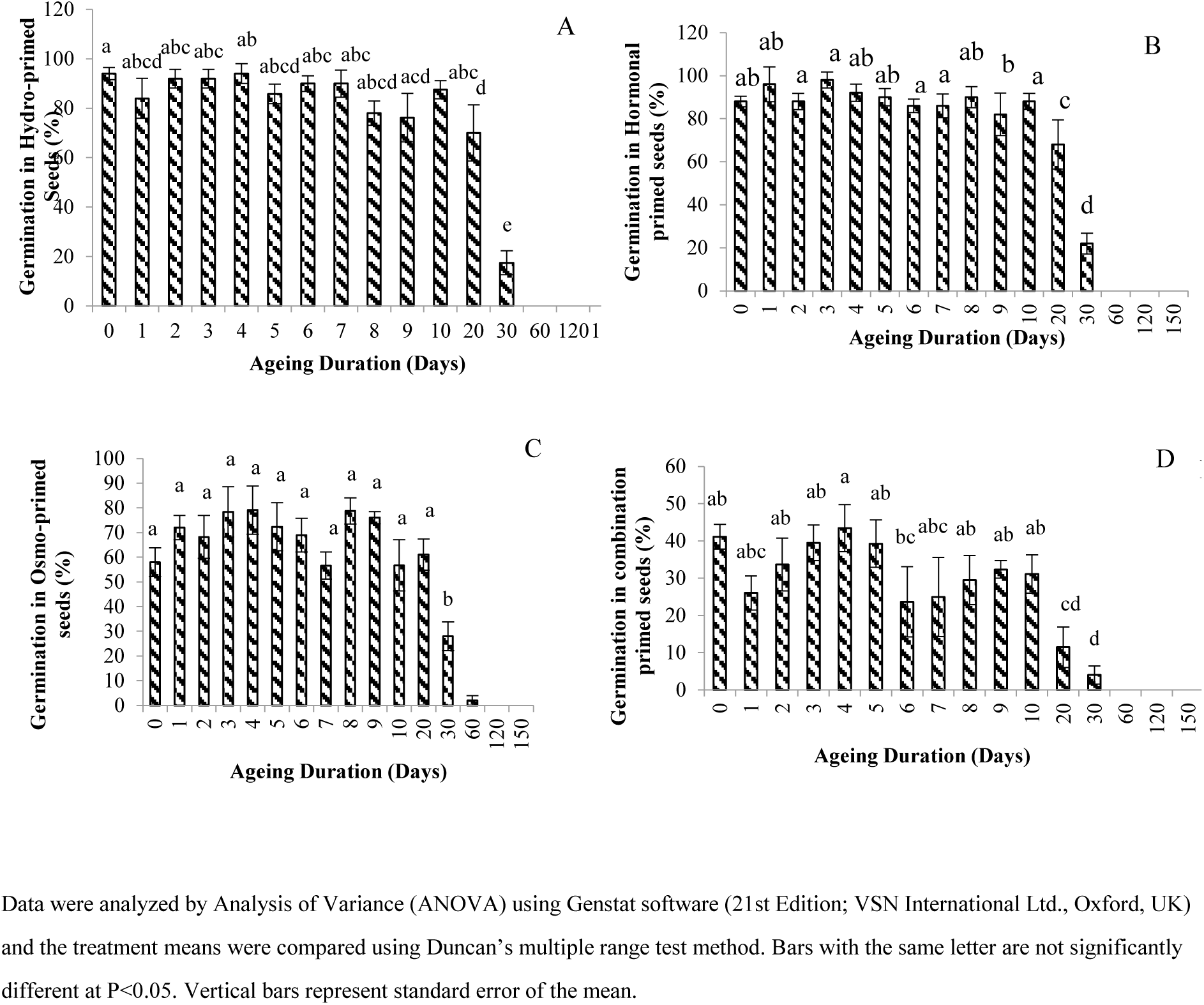
Germination percent in hydro (A), hormone (B), osmo (C) and combination-primed (D) *Vachellia pachyceras* seeds after longevity studies.

**Table 6.** Effect of Seed Priming Treatments on Germination Parameters of *Vachellia pachyceras* Seeds. Treatment-Hydropriming.

| **Treatment- Hydropriming** |  |  |  |
|---|---|---|---|
| Ageing Duration (Days) | Germination Rate (Days) | Seedling Vigor Index (SVI) | Time to 50% germination (Days) |
| 0 (Control) | 66.3 ± 2 abc | 0.233 ± 0.02 a | 1.38 ± 0.1 ab |
| 1 | 61.2 ± 6 bc | 0.159 ± 0.02 bcd | 1.25 ± 0.1 abc |
| 2 | 78.9 ± 7 a | 0.235 ± 0.05 a | 1.35 ± 0.1 ab |
| 3 | 65.1 ± 5 abc | 0.179 ± 0.01 abc | 1.35 ± 0.1 ab |
| 4 | 58.8 ± 9 bc | 0.182 ± 0.02 abc | 1.41 ± 0.1 a |
| 5 | 59.9 ± 2 bc | 0.154 ± 0.01 bcd | 1.22 ± 0.1 abc |
| 6 | 73.8 ± 9 ab | 0.201 ± 0.03 ab | 1.29 ± 0.1 abc |
| 7 | 49.3 ± 1 c | 0.132 ± 0.01 cde | 1.33 ± 0.1 ab |
| 8 | 57.6 ± 3 bc | 0.117 ± 0.03 de | 1.09 ± 0.1 bc |
| 9 | 49.1 ± 6 c | 0.086 ± 0.01 ef | 1.20 ± 0.1 abc |
| 10 | 52.9 ± 3 c | 0.124 ± 0.01 cde | 1.25 ± 0.1 abc |
| 20 | 28.1 ± 3 d | 0.058 ± 0.01 fg | 1.01 ± 0.1 c |
| 30 | 27.4 ± 4 d | 0.017 ± 0.01 g | 0.42 ± 0.1 d |
| 60 | - | - | - |
| **Treatment- Hormonal priming** |  |  |  |
| Ageing Duration (Days) | Germination Rate | Seedling Vigor Index (SVI) | Time to 50% germination (Days) |
| 0 (Control) | 74.5 ± 7 abc | 0.202 ± 0.03 abc | 0.78 ± 0.1 cd |
| 1 | 75.1 ± 8 ab | 0.231 ± 0.03 ab | 0.92 ± 0.2 cd |
| 2 | 87.9 ± 2 a | 0.248 ± 0 02 abc. | 0.58 ± 0.0 d |
| 3 | 73.0 ± 5 bcd | 0.234 ± 0.04 abc | 0.63 ± 0.2 d |
| 4 | 75.9 ± 7 abc | 0.217 ± 0.07 abc | 0.76 ± 0.1 cd |
| 5 | 63.8 ± 5 cde | 0.278 ± 0.04 a | 1.13 ± 0.1 cd |
| 6 | 60.0 ± 2 def | 0.201 ± 0.05 abc | 1.00 ± 0.1 cd |
| 7 | 57.0 ± 7 ef | 0.171 ± 0.05 abcd | 1.29 ± 0.2 bcd |
| 8 | 58.0 ± 4 ef | 0.21 ± 0.03 abc | 1.16 ± 0. 2 cd |
| 9 | 48.4 ± 6 f | 0.144 ± 0.02 cd | 1.38 ± 0.1 bcd |
| 10 | 47.2 ± 3 f | 0.158 ± 0.02 bcd | 1.47 ± 0. 2 bc |
| 20 | 29.0 ± 4 g | 0.068 ± 0.01 de | 1.99 ± 0. 1 b |
| 30 | 18.8 ± 2 g | 0.016 ± 0.00e | 4.65 ± 0.7 a |
| 60 | - | - | - |
| **Treatment- Osmopriming** |  |  |  |
| Ageing Duration (Days) | Germination Rate | Seedling Vigor Index (SVI) | Time to 50% germination (Days) |
| 0 (Control) | 50.7 ± 5 abc | 0.094 ± 0.02 abc | 1.53 ± 0.0 cde |
| 1 | 55.9 ± 5 a | 0.137 ± 0.02 a | 1.52 ± 0.2 cde |
| 2 | 43.6 ± 3 bcd | 0.085 ± 0.02 bc | 1.62 ± 0.1 cde |
| 3 | 50.4 ± 3 abc | 0.124 ± 0.02 ab | 1.32 ± 0.2 de |
| 4 | 53.4 ± 4 ab | 0.134 ± 0.02 a | 1.18 ± 0.0 e |
| 5 | 33.7 ± 1 d | 0.077 ± 0.01 bc | 1.90 ± 0.2 cde |
| 6 | 38.4 ± 3 d | 0.08 ± 0.01 bc | 1.75 ± 0.2 cde |
| 7 | 39.2 ± 4d | 0.072 ± 0.02 cde | 1.64 ± 0.1 cde |
| 8 | 39.2 ± 4 cd | 0.102 ± 0.02 abc | 1.83 ± 0.2 de |
| 9 | 35.1 ± 3 d | 0.081 ± 0.01 bc | 2.14 ± 0.3 c |
| 10 | 33.9 ± 3 d | 0.074 ± 0.02 bcd | 1.73 ± 0.0 cde |
| 20 | 40.0 ± 7 cd | 0.070 ± 0.03 cde | 2.02 ± 0.4 cd |
| 30 | 20.8 ± 2 e | 0.020 ± 0.01 d | 4.25 ± 0.5 b |
| 60 | 11.1 ± 0 e | 0 | 8.5 ± 0.0 a |
| **Treatment- Combination priming** |  |  |  |
| Ageing Duration (Days) | Germination Rate | Seedling Vigor Index (SVI) | Time to 50% germination (Days) |
| 0 (Control) | 54.3 ± 1 a | 0.122 ±0.05 a | 1.33 ± 0.3 d |
| 1 | 55.2 ±1 a | 0.059 ± 0.02 b | 1.40 ± 0.3 d |
| 2 | 33.5 ± 2 ab | 0.048 ± 0.01 b | 2.65 ± 0.3 c |
| 3 | 36.4 ± 5 ab | 0.051 ± 0.01 b | 2.27 ± 0.3 cd |
| 4 | 52.1 ± 9 a | 0.083 ± 0.01 ab | 1.63 ± 0.4 cd |
| 5 | 39.5 ± 4 ab | 0.055 ± 0.01 b | 2.00 ± 0.3 cd |
| 6 | 43.6 ± 7 a | 0.051 ± 0.02 b | 1.92 ± 0.4 cd |
| 7 | 37.2 ± 5 ab | 0.040 ± 0.01 b | 1.89 ± 0.2 cd |
| 8 | 39.5 ± 3 ab | 0.042 ± 0.01 b | 1.97 ± 0.2 cd |
| 9 | 33 ± 4 ab | 0.032 ± 0.00 b | 2.35 ± 0.2 cd |
| 10 | 39.9 ± 5 ab | 0.047 ± 0.01 b | 1.63 ± 0.2 cd |
| 20 | 17.2 ± 4 b | 0.010 ± 0.00 b | 4.75 ± 1.1 b |
| 30 | 13.3 ± 3 b | - | 6.75 ± 1.3 a |
| 60 | - | - | - |

Data were analyzed by Analysis of Variance (ANOVA) using Genstat software (21st Edition; VSN International Ltd., Oxford, UK) and the treatment means were compared using Duncan’s multiple range test method. Mean values with the same letter are not significantly different at P<0.05.

Hormone primed *V. pachyceras* seeds, artificially aged for various durations, exhibited variation in germination parameters. Seeds aged for two days had the highest germination rate (Table 6), and seeds aged for three days had the highest germination percentage (Figure 2B), and both the parameters declined as the ageing duration increased (Figure 2B & Table 6). However, time to 50% germination (days) was the lowest for 2-day aged seeds but increased thereafter for seeds that were aged up to 30 days (Table 6). The SVI was the highest at 5-days of ageing and declined steadily up to 30 days.

Osmoprimed *V. pachyceras* seeds aged for 1 day had the highest germination rate, and no germination was observed after 60 days of ageing (Table 6; Figure 2C). The SVI was the highest for seeds that were aged for one day and declined after 4 days (Table 6). Germination percentage remained significantly higher for seeds aged between 0 days and 20 days but declined sharply thereafter (Figure 2C). Time to 50% germination was the shortest for seeds aged for 4 days and continued to increase (number of days) progressively with longer ageing duration (i.e., up to 60 days) (Table 6).

Similar results were observed for different germination parameters when *V. pachyceras* seeds were primed with combination (both hydropriming and osmopriming treatments) treatments and subjected to different ageing durations. For example, germination rate was the highest for seeds aged for 0 or 1 day and continued to decline as the ageing duration increased up to 30 days (Table 6). Similarly, SVI was the highest for seeds that were aged 0 day and continued to decline up to 20 days (Table 6). Germination percentage was the highest for seeds at 4 days of ageing and declined as the ageing duration increased up to 30 days (Figure 2D). The shortest duration (number of days) to achieve 50% germination was observed in seeds that were not aged or aged for 1 day and it increased as the ageing duration increased (Table 6).

### C. oxyphyllus

Data of probit analysis and subsequent ANOVA revealed that hydro-priming (F _3, 19_ = 37.62, *p* < .001) resulted in significantly higher p50 values (i.e., number of days to lose seed viability to 50%; i.e., 74 days) than the hormone primed, osmoprimed, and combination primed *C. oxyphyllus* seeds. No significant difference in p50 values were detected between hormone primed, osmoprimed, and combination primed *C. oxyphyllus* seeds (Figure 3 and Table 7).

**Figure 3.**
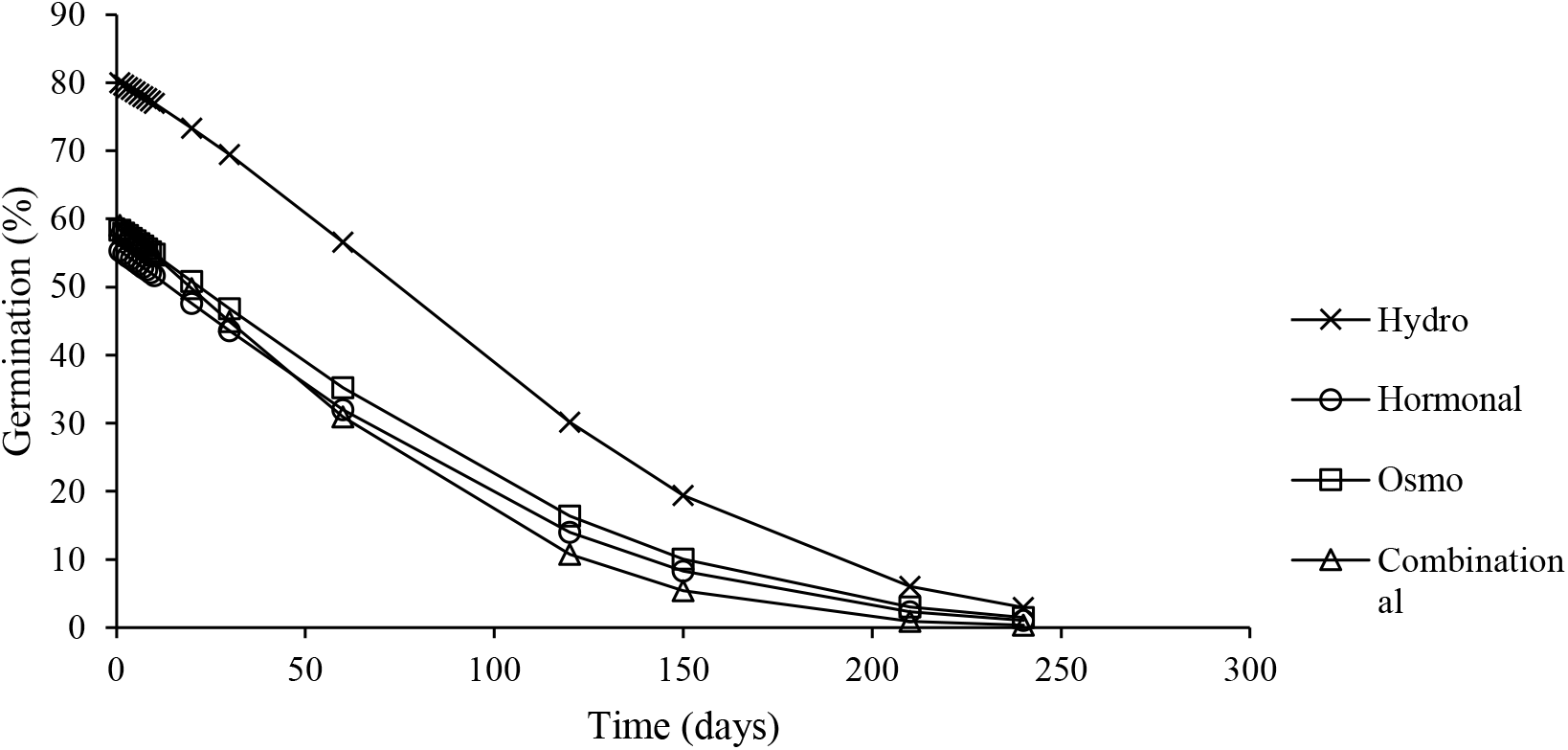
Survival curves fitted using a generalized linear model with binomial error and probit link function using ®GenStat 21^st^ Edition (VSN International Ltd., Oxford, UK) for ageing experiments in hydro-primed, hormone primed, osmoprimed, and combination (hydro + osmo) primed *Convolvulus oxyphyllus* seeds.

**Table 7.** Effect of Seed Priming Treatments on Germination Parameters of *C. oxyphyllus* seeds. Treatment-Hydropriming.

| **Treatment- Hydropriming** |  |  |  |
| Ageing Duration (Days) | Germination Rate | Seedling Vigor Index (SVI) | Time to 50% germination (Days) |
|---|---|---|---|
| 0 (Control) | 19.1 ± 2 bcde | 0.002 ± 0.001 dfg | 3.28 ± 0.9 bcde |
| 1 | 16.6 ± 3 de | 0.001 ± 0.000 g | 2.25 ± 0.1 de |
| 2 | 18.8 ± 3 bcde | 0.001 ± 0.001 fg | 3.83 ± 0.8 bcd |
| 3 | 36.2 ± 5 a | 0.002 ± 0.001 dfg | 1.97 ± 0.1 e |
| 4 | 23.1 ± 4 ab | 0.006 ± 0.003 bcde | 1.70 ± 0.4 e |
| 5 | 25.9 ± 5 abcde | 0.011 ± 0.002 a | 2.33 ± 0.1 cde |
| 6 | 33.3 ± 8 ab | 0.008 ± 0.001 abc | 1.79 ± 0.2 c |
| 7 | 27.5 ± 8 abcd | 0.008 ± 0.002 ab | 2.73 ± 0.6 cde |
| 8 | 31.4 ± 4 abcd | 0.002 ± 0.000 defg | 2.15 ± 0.3 de |
| 9 | 29.9 ± 6 abcd | 0.001 ± 0.000 g | 2.53 ± 0.5 cde |
| 10 | 25.7 ± 4 abcde | 0.003 ± 0.001 dfg | 2.88 ± 0.3 cde |
| 20 | 34.0 ± 3 a | 0.007 ± 0.000 abcd | 2.30 ± 0.3 de |
| 30 | 30.8 ± 3 abcd | 0.004 ± 0.000 cdef | 2.69 ± 0.3 cde |
| 60 | 32.6 ± 3 abc | 0.001 ± 0.000 g | 2.48 ± 0.2 cde |
| 120 | 12.2 ± 2 e | 0.001 ± 0.000 g | 7.35 ± 1.1 a |
| 150 | 16.9 ± 3 cde | 0.001 ± 0.000 g | 4.25 ± 1.5 bc |
| 210 | 20.0 ± 0 * | 0.000 ± 0.000 * | 5.50 ± 0.0 * |
| 240 | - | - | - |
| **Treatment- Hormone-priming** |  |  |  |
| Ageing Duration (Days) | Germination Rate | Seedling Vigor Index (SVI) | Time to 50% germination (Days) |
| 0 (Control) | 45.4 ± 5 ab | 0.0067 ± 0.001a | 1.96 ± 0.0 e |
| 1 | 37.5 ± 5 b | 0.0028 ± 0.001 cd | 1.90 ±0.2 e |
| 2 | 39.9 ± 6 ab | 0.0049 ± 0.001abc | 1.68 ± 0.1 e |
| 3 | 42.0 ± 3 ab | 0.0051 ± 0.001 abc | 1.92 ± 0.2 e |
| 4 | 35.9 ± 6 b | 0.0046 ± 0.001 abc | 1.61 ± 0.1 e |
| 5 | 38.7 ± 4 ab | 0.0054 ± 0.001 a | 1.80 ± 0.2 e |
| 6 | 45.6 ± 1 ab | 0.0031 ± 0.001 bcd | 1.63 ± 0.0 e |
| 7 | 39.1 ± 4 ab | 0.0026 ± 0.001 cd | 1.97 ± 0.2 e |
| 8 | 34.6 ± 6 bc | 0.0030 ±0.000 cd | 3.02 ± 0.9 d |
| 9 | 39.9 ± 2 ab | 0.0028 ± 0.001 cd | 1.83 ± 0.1 e |
| 10 | 44.4 ± 2 ab | 0.0025 ± 0.001 cd | 1.65 ± 0.1 e |
| 20 | 44.7 ± 3 ab | 0.0022 ± 0.001 bcd | 1.76 ± 0. 2 e |
| 30 | 45.5 ± 2 ab | 0.0009 ± 0.000 d | 1.68 ± 0.1 e |
| 60 | 22.0 ± 9 cd | 0.0009 ± 0.000 d | 2.63 ± 0.7 de |
| 120 | 54.2 ± 4 * | 0.0019 ± 0.000 bcd | 1.38 ± 0.1 e* |
| 150 | 4.8 ± 3 * | 0.0002 ± 0.000 d | 5.00 ± 2.0 c* |
| 210 | 11.1 ± 0 * | 0.0001 ± 0.000 d | 8.50 ± 0.0 b* |
| 240 | 8.8 ± 1 * | 0.0001 ± 0.000 d | 11.00 ± 1.50 a* |
| **Treatment- Osmopriming** |  |  |  |
| Ageing Duration (Days) | Germination Rate | Seedling Vigor Index (SVI) | Time to 50% germination (Days) |
| 0 (Control) | 34.8 ± 3 abc | 0.0018 ± 0.0004 bc | 2.26 ± 0.2 acde |
| 1 | 35.8 ± 2 abcd | 0.0034 ± 0.0007 a | 2.07 ± 0.2 de |
| 2 | 39.9 ± 3 a | 0.0022 ± 0.0004 bc | 2.06 ± 0.2 de |
| 3 | 35.7 ± 3 abcd | 0.0025 ± 0.0005 ab | 2.03 ± 0.3 de |
| 4 | 26.7 ± 2 bcd | 0.0022 ± 0.0002 bc | 2.75 ± 0.3 abcd |
| 5 | 29.7 ± 2 abcd | 0.0017 ± 0.0003 b | 2.3 ± 0.3 acde |
| 6 | 36.4 ± 2 abc | 0.0018 ± 0.0002 bc | 2.25 ± 0.1 acde |
| 7 | 37.3 ± 2ab | 0.0017 ± 0.0001 bc | 2.20 ± 0.3 acde |
| 8 | 29.7 ± 3 abcd | 0.0018 ± 0.0001 bc | 2.87 ± 0.3 abcd |
| 9 | 36.3 ± 2 abc | 0.0018 ± 0.0005 bc | 2.23 ± 0.1 acde |
| 10 | 29.5 ± 4.abcd | 0.0018 ± 0.0002 bc | 3.05 ± 0.9 abcd |
| 20 | 34.7 ± 3 abcd | 0.0019 ± 0.0002 bc | 2.43 ± 0.3 acde |
| 30 | 24.4 ± 0 d | 0.0000 ± 0.0000 c | 3.57 ± 0.1 ab |
| 60 | 27.5 ± 3 bcd | 0.0003 ± 0.0000 c | 3.15 ± 0.4 abc |
| 120 | 50.0 ± 0 * | 0.0001 ± 0.0000 c | 1.50 ± 0.0 * |
| 150 | 16.7 ± 0 * | - | 4.00 ± 0.0 * |
| 210 | 62.5 ± 12 * | - | 1.25 ± 0.3 * |
| 240 | - | - | - |
| **Treatment- Combination priming** |  |  |  |
| Ageing Duration (Days) | Germination Rate | Seedling Vigor Index (SVI) | Time to 50% germination (Days) |
| 0 (Control) | 44.0 ± 3.3 abc | 0.0033 ± 0.0004 b | 1.74 ± 0.1 b |
| 1 | 34.6 ± 1.8 cdef | 0.005 ± 0.0010 a | 2.22 ± 0.1 b |
| 2 | 49.8 ± 4.7 a | 0.003 ± 0.0005 bcd | 2.17 ± 0.4 b |
| 3 | 35.4 ± 2.0 cdef | 0.0015 ± 0.0002 cdefg | 2.33 ± 0.3 b |
| 4 | 42.3 ± 1.1 abcde | 0.0015 ± 0.0002 cdefg | 1.76 ± 0.0 b |
| 5 | 37.2 ± 4.4 cdef | 0.0020 ± 0.0003 bcd | 2.25 ± 0.6 b |
| 6 | 46.8 ± 2.0 ab | 0.0027 ± 0.0004 bcde | 1.63 ± 0.1 b |
| 7 | 32.7 ± 1.1 dfg | 0.0019 ± 0.0007 bcdefg | 1.98 ± 0.1 b |
| 8 | 38.6 ± 1.6 bcdef | 0.0021 ± 0.0003 bcdefg | 2.02 ± 0.1 b |
| 9 | 40.5 ± 1.0 bcdef | 0.0022 ± 0.0007 bcdefg | 1.97 ± 0.1 b |
| 10 | 37.9 ± 1.7 bcdef | 0.0023 ± 0.0005 bcdef | 1.77 ± 0.1 b |
| 20 | 38.2 ± 4.1 bcdef | 0.0032 ± 0.0000 abc | 1.91 ± 0.2 b |
| 30 | 25.7 ± 4.1 g | 0.0005 ± 0.0002 cefg | 2.4 ± 0.2 b |
| 60 | 42.5 ± 3.8 abcd | 0.0002 ± 0.0001 ceg | 2.17 ± 0.1 b |
| 120 | - | - | - |
| 150 | 7.7 ± 0.6 * | - | 12.5 ± 1.0 a* |
| 210 | - | - | - |
| 240 | - | - | - |

Data were analyzed by Analysis of Variance (ANOVA) using Genstat software (21st Edition; VSN International Ltd., Oxford, UK) and the treatment means were compared using Duncan’s multiple range test method. Bars with the same letter are not significantly different at P<0.05. Vertical bars represent standard error of the mean.
\*Late ageing metrics are dominated by sampling noise.

Ageing duration (number of days), in general, affected the germination parameters of hydroprimed *C. oxyphyllus* seeds. Increased ageing duration reduced germination percentage (Figure 4A), germination rate, SVI, and increased time to 50% germination (Table 7). For example, germination rate was the highest for seeds aged for three days and declined thereafter. However, unlike *V. pachyceras* seeds, *C. oxyphyllus* seeds, were artificially aged up to 210 days, continued to germinate successfully (Table 7), although germination percentage declined beyond 210 days (Figure 4A).

**Figure 4.**
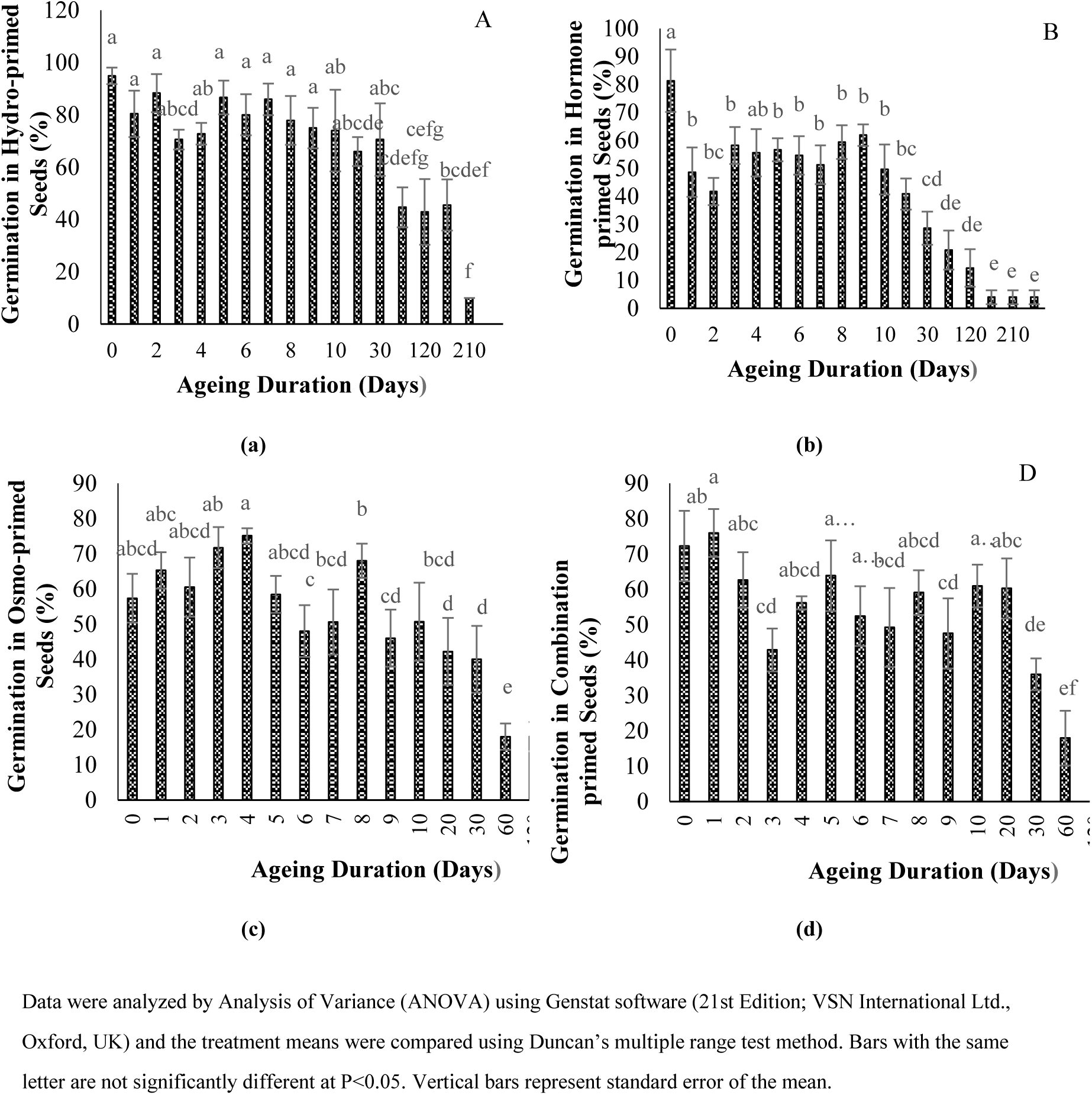
Germination parameters in hydro (A), hormone (B), osmo (C) and combination-primed (D) *Convolvulus oxyphyllus* seeds after longevity studies.

The SVI was the highest for hydroprimed *C. oxyphyllus* seeds, aged for five days, declined up to 10 days and then slightly increased but continued to decrease again until 210 days (Table 7). The highest germination percentage was observed in seeds aged for 0, 1, 2, 5, 6, 7, and 8 days but declined when the ageing duration increased (Figure 4A). The shortest time to reach 50% germination was achieved in seeds aged for either 3, 4, or 6 days; however, it increased as the ageing duration (days) increased (Table 7).

Hormone (GA_3_) primed *C. oxyphyllus* seeds, when aged artificially at various durations, affected different germination parameters. Seeds aged for 120 days had the highest germination rate (Table 7), which declined thereafter as the ageing duration increased up to 240 days. The SVI was the highest for 0-day aged seeds and declined steadily up to 240 days of ageing (Table 7). Similarly, germination percentage was the highest for seeds that were aged for 0 day and continued to decline up to 240 days of ageing (Figure 4B). Time to 50% germination (days) was the lowest for seeds that were aged between 0 day and 120 days, with the exception of those aged for 8 and 60 days, and steadily increased thereafter up to 240 days (Table 7).

Osmoprimed *C. oxyphyllus* seeds aged for 210 days exhibited the highest germination rate, whereas no seed germination was observed for seeds aged for 240 days (Table 7). The SVI was the highest for seeds aged for 1 day and declined thereafter. SVI could not be determined for seeds that were aged more than 120 days due to high mortality after germination leading to absence of data for biomass (Table 7). Similarly, germination percentage remained significantly high for seeds that were aged between 0 day and 4 days but declined considerably for seeds that were aged beyond 30 days (Figure 4C). Seeds that were aged for 210 days required the shortest time (1.25 days) to reach 50% germination, which was significantly lower than other ageing treatments, however, it should be noted that the total number of seeds germinated was also low for 210 days (Table 7).

Combination primed seeds showed similar patterns with varying magnitudes across ageing treatments (Figure 4D and Table 7). For instance, the germination rate was the highest for seeds aged for two days (Table 7). For seeds aged between 3 and 60 days, the germination rate was still relatively high but no germination was observed for seeds that were aged for 120, 210, and 240 days (Table 7). Similarly, the SVI was the highest for seeds that were aged for 1 day. The SVI declined for seeds that were aged for 2 to 4 days but started to increase again for seeds that were aged between 5 days and 20 days, which drastically declined again (Table 7). Germination percentage was the highest for seeds that were aged for 1 day and declined as the ageing duration increased up to 150 days (Figure 4D). The shortest time to achieve 50% germination was observed with seeds that were aged for either 0 to 60 days and increased as the ageing duration increased up to 150 days (Table 7).

## 4. Discussion

Seedling survival after emergence is a critical determinant of restoration success especially in arid ecosystems, where narrow and unpredictable rainfall windows severely restrict opportunities for establishment. As the natural recruitment of seedlings is limited by these factors, restoration efforts depend on maximizing germination and emergence during the short favorable period. Seed enhancement techniques like priming offers a promising approach for accelerating and synchronizing germination, thereby increasing the probability of seedling survival. Though there are benefits for priming, its effect on seed longevity in the native species is yet to be explored. The current study provides a systematic assessment of germination responses and artificial ageing trajectories in the primed seeds to choose the suitable priming treatment for *V. pachyceras* and *C. oxyphyllus* enabling the identification of priming treatments that simultaneously optimize germination and preserve seed longevity for use in large-scale restoration programs.

In this study, seeds of two native plant species were subjected to priming to trigger the germination process and artificially age them at 45 °C for different durations to evaluate the decline in their viability. The time taken by 50% seeds to lose their viability (p50), estimated by probit analysis, is often used as a measure of relative seed longevity [34,61]. As a measure of seed longevity across species, the p50 value has been shown to vary in relation to seed structure and microclimate at the geographic origin of the species [27]. While Probert et al., [27] reported estimates of p50 ranging from 0.1 to 771 days, and Mondoni et al., [34] classified seed longevity as following: (a) p50 ≤ 1 d indicates ‘very short’ seed longevity; (b) p50 > 1 to ≤ 10, ‘short’; (c) p50 > 10 to ≤ 100, ‘medium’; (d) p50 > 100 to ≤ 1000 ‘long’; and (e) p50 > 1000, ‘very long’.

As per the classification of Mondoni et al. [34] the p50 value of hormone primed (22 days) and hydroprimed (21 days) *V. pachyceras* seeds from this study can be termed as ‘medium’ longevity. This is consistent with the findings of Suleiman et al. [43], where only mechanically scarified *V. pachyceras* seeds subjected to ageing (at 45 °C) had a p50 value of 39 days, which is also categorized as ‘medium’ longevity. The comparatively lower p50 value in this study may be attributed to the priming treatment applied after mechanical scarification, which accelerated the germination process. Such hydration induced metabolic activities has been reported to have negative influence on storage duration as suggested by Fabrissin et al. [29].

It is evident that the effect of priming treatments and its response in the longevity of the seed is species specific and priming method specific, reflecting intrinsic differences in seed structure, and tolerance to hydration – dehydration cycles. In *V. pachyceras*, both hydro and hormonal treatments resulted in similar p50 values with no significant difference. The absence of significant differences in p50 value indicates that both priming treatments induced similar level of controlled metabolic activation, with minimal post-hydration deterioration and provided comparable protection against seed deterioration during storage. Additionally, this response was consistent with models proposed by Ellis and Roberts [59], which suggest mild hydration-dehydration cycles can promote membrane reorganization without triggering rapid deterioration. From these results, it can be suggested that controlled hydration (Hydropriming) alone appears sufficient to achieve good germination percentage and longevity.

It is also important to note that there was no significant difference in the germination percentage in hormonal and hydroprimed seeds, aged up to 10 days. As discussed earlier, priming initiates cell metabolic activity consuming the starch reserves [62]. When seeds are redried before radicle emergence, the seed enters a state of physiological readiness. This may render them metabolically vulnerable due to reduced metabolic integrity and depleted anti-oxidant levels. Seed viability decline gradually as the activities of damage repairing enzymes decrease resulting in unrepaired oxidative damage [63] The lack of significant decline in germination percentage of hydro and hormonal primed seeds of *V. pachyceras* aged artificially for up to 10 d suggests that physiological and biochemical integrity of the seeds was largely maintained during this period.

This supports the conclusion that 10 days of artificial ageing is physiologically safe to store hormonal or hydroprimed *V. pachyceras* seeds. Results of studies by Sharma et al. [64] also indicated positive impact of priming on longevity of *Vigna radiata* from Fabaceae family after 8 days of artificial ageing at 40 °C and 90% RH.

In contrast, osmopriming accelerated the viability loss possibly due to the osmotic stress affecting ion homeostasis and thus resulting in faster deterioration of seeds during subsequent storage of this species The prolonged exposure to low water potential during osmopriming can induce oxidative damage and reduce the seed quality [65]. However, this finding contrasts with the findings of Ren et al. [66], which suggested slower deterioration in osmoprimed rice seeds compared to hydroprimed seeds under accelerated ageing reinforcing the species specific nature of priming responses.

The ineffectiveness of combination priming in *V. pachyceras* suggests that increasing priming intensity or using complex procedures might disrupt seed physiology and negatively affect the seed quality rather than enhancing.

Hydropriming resulted in significantly higher p50 value (74 days) for *C. oxyphyllus* seeds compared to hormone priming, osmopriming, and combination priming treatments, and, therefore, can be classified as ‘medium’ longevity. As all the other priming treatments shortened the longevity of *C. oxyphyllus*, it appears that only hydropriming provided sufficient metabolic activation without causing deterioration of seed viability in this species. The effectiveness of hydropriming depends on initial seed viability, genotype, environmental adaptation and ability to re-establish desiccation tolerance after priming [29]. Though a declining trend was noticed in seeds aged 0-30 d, there was no significant difference in the germination percentage. Therefore, this duration may be considered a safe storage period of hydroprimed *C. oxyphyllus* without any considerable change in seed quality.

The difference in response of these two species to various priming treatments confirms a strong link between seed structural traits, ecological origin, and storage capability [27]. Though, artificial ageing p50 values provide us a comparative method to measure the longevity among the priming treatments, it cannot be converted into real time storage durations. As the species specific constants [59] for *V. pachyceras* and *C. oxyphyllus* are not available, the p50 estimates should be considered as relative indicators of storage rather than the real time shelf life of treated seeds.

Additionally, literature on the effect of hydropriming, hormone priming, osmopriming or combination priming on seed longevity of desert shrub species is not currently available. In the absence of comparable information, the findings of this study on the effects of different kinds of priming can be used as a guideline to estimate the seed longevity of different native species in Kuwait and surrounding regions, where environmental conditions and native species distribution are similar. In the longevity studies, due to the scarcity of the native seeds, and limited resources, one batch of seeds were retrieved in the predetermined times divided into subsamples and considered as replicates to conduct the experiments. This might have resulted to overestimated degrees of freedom. Hence, the interpretation of the results should be made with caution and future studies should include multiple independently aged batches at each ageing interval to permit valid estimation of between batch variability and appropriate mixed effect analysis.

The current study addressed several important research gaps identified by Hasanovic et al. [25] in priming treatment and its effect on longevity. Specifically, it determined the threshold of imbibition duration required to avoid premature germination and standardized species specific priming protocols for *V. pachyceras* and *C. oxyphyllus.* However, this study represents only the initial step toward addressing the broader knowledge gap. Further investigations are required to elucidate physiological and biochemical changes associated with seed priming and ageing in native plants. In addition, extending the research to other ecologically and economically important native plants is essential for developing scalable seed enhancement techniques and promoting their wider adoption within commercial seed industry.

From a broader perspective, efficient and effective utilization of limited native plant seeds remains a strong viable option for large-scale revegetation and restoration activities particularly because seedling production and outplanting are generally more expensive and labor intensive than direct seeding. The findings of the study demonstrate that species-specific seed enhancement techniques can promote better and uniform germination, thereby enhancing seedling establishment and increasing the overall success and cost effectiveness of revegetation programs [67].

To conserve and expand the native plant cover in the desert ecosystems of Kuwait and other arid regions, systematic and rigorous studies are required to develop cost-effective and easily adaptable seed technologies. Such efforts will facilitate the implementation of best practices for seed germination and seedling establishment ultimately supporting the conservation of native plant species, and improving the success of seed based restoration and revegetation activities.

## 5. Conclusions

The study ranks longevity among the priming treatments for *Vachellia pachyceras* and *Convolvulus oxyphyllus* and identifies that hydropriming is the least detrimental for both species. Though hormonal priming in *V. pachyceras* resulted in slightly superior germination percentage among the priming treatments, it was not significantly higher than the hydropriming. Additionally, hydropriming is recognized as a more efficient, cost-effective, and rapid method, as it involves simple soaking in water rather than complex procedures. Therefore, hydropriming is recommended for both species considering the ease of execution by ecologists for practical application in terms of higher germination, and due to its simplicity and economic benefits.

## Institutional Review Board Statement

Not applicable.

## Informed Consent Statement

Not applicable.

## Data Availability Statement

Data are contained in the text, tables and figures.

## Acknowledgements

The authors would like to express their sincere appreciation to the management of Kuwait Institute for Scientific Research (KISR) and Kuwait Foundation for the Advancement of Sciences (KFAS) Kuwait, for its continued interest, encouragement, logistical and financial support. The authors extend their gratitude to Dr. Andrea Mondoni for his valuable suggestions and guidance to perform probit analysis, and Mr. Ijas Mohamed Rafi, ELSRC-KISR in preparing the supplementary graphical representation of priming procedures.

